# Clustering instantaneous phase-locking states with a rank-two Bures-Wasserstein distance

**DOI:** 10.64898/2026.11.03.764018

**Authors:** Maliheh Miri, Harry H. Behjat, Dimitri Van De Ville, Anders S. Olsen

## Abstract

Instantaneous phase-locking (iPL) matrices, defined by the cosines of pairwise phase differences between band-limited signals, underpin widely used methods for time-resolved functional connectivity, including Leading Eigenvector Dynamics Analysis (LEiDA). Each iPL matrix has rank at most two and is fully characterized by its two nonzero eigenvalues and corresponding eigenvectors. Yet, existing methods retain different parts of this decomposition and compare them using distances that do not reflect the geometry of the underlying sample space. We show that weighting the two eigenvectors by the square roots of their eigenvalues recovers each channel’s phase relative to the mean phase, and that optimal orthogonal Procrustes alignment of these weighted eigenvector pairs yields the Bures–Wasserstein distance between the corresponding iPL matrices. In EEG, where most channels oscillate nearly in phase, existing distances become insensitive to phase-lag size, whereas the Bures–Wasserstein distance preserves this sensitivity. In simulations, clustering with this distance recovered states differing in both lag size and spatial pattern that existing distances failed to distinguish. In resting-state EEG, it resolved distinct phase-gradient states that other methods merged into a single global in-phase state.

## 1. INTRODUCTION

Brain connectivity patterns vary over time, and dynamic functional connectivity (dFC) methods go beyond the summarized view of static connectivity [1]. One class of dFC methods derives connectivity from instantaneous co-activation, providing maximal temporal resolution [2]. In EEG, brain states are most often studied with microstate analysis, which clusters instantaneous scalp-potential topographies into typically four classes [3]. Phase-locking methods instead characterize the relative timing of oscillations across channels [4].

Leading Eigenvector Dynamics Analysis (LEiDA) [2] computes the instantaneous phase-locking (iPL) matrix of pairwise cosine phase differences at each time point, retains only its leading eigenvector, and clusters these vectors into recurring states. The leading eigenvector was motivated to reduce noise and dimensionality. However, the iPL matrix has rank at most two, and its second eigenvector carries substantial, structured variance; for typical EEG data about a quarter of the matrix trace. Moreover, because eigenvectors are only defined up to sign, LEiDA flips each eigenvector to a fixed sign convention before *K*-means. This restricts the data to one half of the sphere along a nonlinear boundary, which can split clusters and is incompatible with the sign-symmetric sample space of eigenvectors [5, 6]. LEiDA has mainly been applied to fMRI [2, 7] where it often yields a dominant “global” in-phase state, often addressed by regressing out the global signal [6].

Eigenvector Dynamics Analysis (EiDA) [8] retains both eigenvectors, but clusters the vectorized upper-triangular iPL matrices using *K*-means and the cosine distance. This treats the matrix entries as an unconstrained Euclidean vector, normalizes away the overall strength of phase locking, and averages to centroids that need not retain the rank-two structure; the distance is thus not derived from the structure of the data. In EEG, the sine of the phase differences is often preferred, as it is insensitive to the zero-lag coupling caused by volume conduction [9].

Here, we ask which distance between the iPL matrices respects their structure. To address eigenvector sign and rotation ambiguities, we align eigenvector pairs using an orthogonal Procrustes transform [10]. We show that when the eigen-vectors are weighted by the square roots of their eigenvalues, the Procrustes distance equals the Bures–Wasserstein (BW) distance between rank-two positive semidefinite matrices [11, 12], an optimal-transport distance. It quantifies the phase shift required to transform one iPL matrix into another, and remains sensitive to the size of phase lags when channels are close to synchrony, as is typical in EEG. We combine it with *K*-means and compare it with alternative distances on simulated and resting-state EEG. Our message is simple: when the real-valued iPL matrix is used, its eigenvalue-weighted eigen-vectors should be compared using the BW distance.

## 2. METHODS

### 2.1. From signals to phase-locking matrices

Let **X** ∈ ℝ^*P×T*^ hold *P* band-pass filtered time series *x*_*j*_ ∈ (*t*) (EEG channels or brain regions), *t* = 1, …, *T*. For each channel, the analytic signal 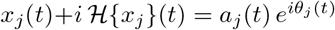 is computed with the Hilbert transform, yielding the instantaneous phase *θ*_*j*_ (*t*), interpretable only for narrow-band signals [13]. For an oscillation that travels across the scalp, phase differences between channels correspond to time lags (Fig. 1a,b); at 10 Hz, 1 rad corresponds to 16 ms. The iPL matrix collects the cosines of all pairwise phase differences [2],

**Fig 1.**
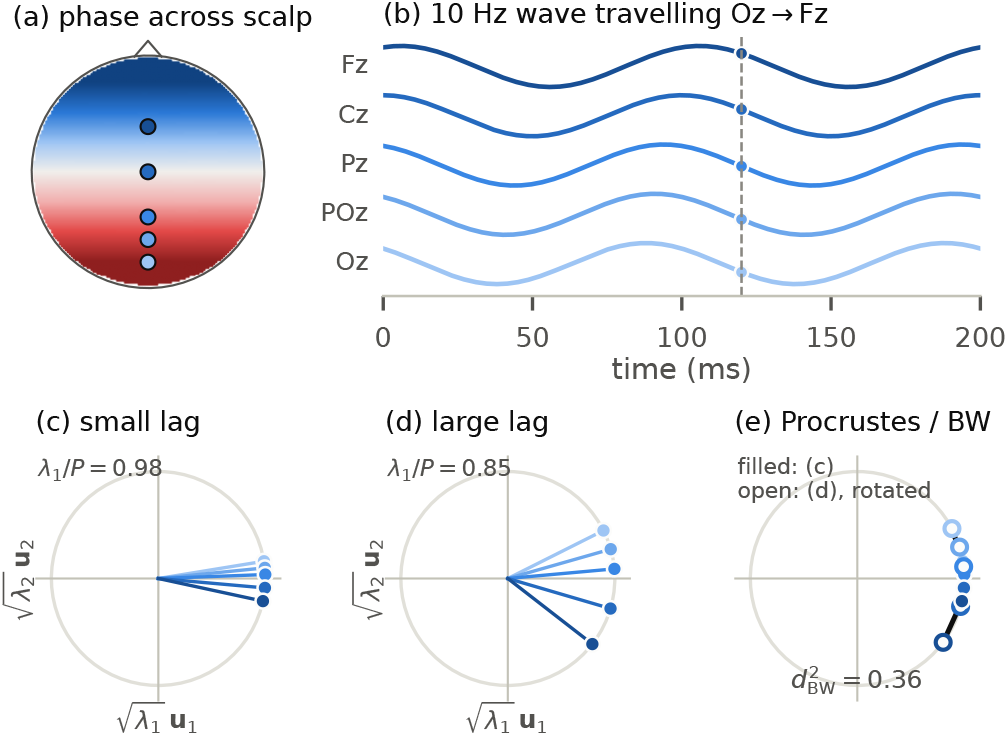
From phases to the BW distance. (a) Relative phase of a 10 Hz wave travelling from occipital to frontal channels (red leads), and (b) signals of five midline channels. (c,d) At one time point, rows of the eigenvalue-weighted eigenvectors **Ũ**_*t*_ are channel phasors relative to the mean phase: a small lag concentrates them (*λ*_1_≈ *P*), a large lag spreads them. (e) The squared BW distance is the summed squared length of the black segments after optimal orthogonal alignment **Ω**.

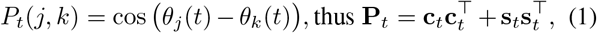

where **c**_*t*_ = cos ***θ***(*t*) and **s**_*t*_ = sin ***θ***(*t*), following from cos(*α*− *β*) = cos *α* cos *β* + sin *α* sin *β* [8]. Hence, **P**_*t*_ has unit diagonal and rank at most two, reducing to rank one when all channel phases coincide modulo *π*. Similar channel phases yield *P*_*t*_(*j, k*)→ 1, while orthogonal phases yield *P*_*t*_(*j, k*)→ 0. Each channel can be represented as a unit-length phasor (cos *θ*_*j*_, sin *θ*_*j*_), and **P**_*t*_ only depends on the angles between the phasors. Consequently, rotating all phasors by a common angle (the ongoing oscillation) or mirroring them (reversing all lags) leaves **P**_*t*_ unchanged.

### 2.2. Eigenvectors and eigenvalues

As **P**_*t*_ has rank at most two, it is fully described by two eigen-vectors **u**_1_, **u**_2_ and eigenvalues *λ*_1_ ≥ *λ*_2_, with *λ*_1_ + *λ*_2_ = *P*.

Collecting the eigenvectors weighted by the square roots of their eigenvalues as 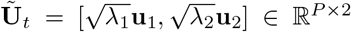, we have

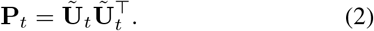

Comparing (2) with (1), the *j*th row of **Ũ**_*t*_ is the phasor of channel *j* relative to the (axial) mean phase of all channels, with cosine and sine components [8] (Fig. 1c,d). Therefore, the eigenvalues quantify the phasor spread: for nearly inphase channels, *λ*_1_≈ *P* and *λ*_2_≈ 0.

This view clarifies the limitations of existing representations. First, the leading eigenvector captures only the cosines; i.e., how far each channel is from the mean phase, not whether it leads or lags. Therefore, two different iPL matrices can share the same leading eigenvector, and near synchrony all leading eigenvectors approach the uniform vector, while the spatial lag pattern resides in the weak second eigenvector. Second, weighting the eigenvectors by the square roots of their eigenvalues restores the unit-length channel phasors. The weighted pair reconstructs the full iPL matrix and determines the sine matrix up to a global sign, 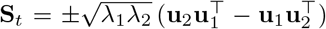. This representation allows comparison through phasor displacements rather than through unweighted eigenvector overlaps.

### 2.3. Existing distances

Clustering requires a distance between the iPL matrices **P**_*a*_ and **P**_*b*_ of two time points. Existing distances (Table 1) can be expressed using the 2 ×2 matrix 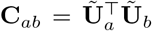. These overlaps are sums of products of cosines and sines of phase differences. Consider two time points with the same spatial lag pattern **g** but different lag size, ***θ***_*a*_ = *ε***g** and ***θ***_*b*_ = (*ε* + *δ*)**g**. Since cos *x* ≈ 1− *x*^2^*/*2 for small *x*, the iPL matrix depends on the lag size only through *ε*^2^, so near synchrony **P**_*a*_ and **P**_*b*_ differ by *O*(*εδ*). Consequently, distances that are smooth functions of **P**_*t*_, its eigenvectors, or their span become insensitive to lag size as the channels approach synchrony. The Grassmann distance further rescales both eigenvectors to unit length, which inflates the weak second direction and its noise, while LEiDA discards the second eigenvector entirely.

**Table 1.**
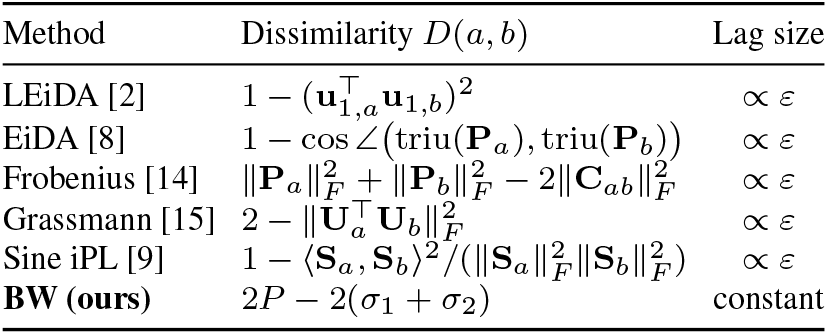
Dissimilarities between iPL matrices **P**_*a*_ and **P**_*b*_, where 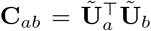 with singular values *σ*_1_, *σ*_2_. Last column: change in distance per unit change in lag size *ε* near synchrony, which vanishes with *ε* for all distances except BW (Sect. 2.3).

| Method | Dissimilarity $D(a,b)$ | Lag size |
|---|---|---|
| LEiDA [2] | $1-(\mathbf{u}_{1,a}^{\top}\mathbf{u}_{1,b})^2$ | ∝ ε |
| EiDA [8] | $1-\cos\angle\big(\mathrm{triu}(\mathbf{P}_a),\mathrm{triu}(\mathbf{P}_b)\big)$ | ∝ ε |
| Frobenius [14] | $\lVert\mathbf{P}_a\rVert_F^2+\lVert\mathbf{P}_b\rVert_F^2-2\lVert\mathbf{C}_{ab}\rVert_F^2$ | ∝ ε |
| Grassmann [15] | $2-\lVert\mathbf{U}_a^{\top}\mathbf{U}_b\rVert_F^2$ | ∝ ε |
| Sine iPL [9] | $1-\langle\mathbf{S}_a,\mathbf{S}_b\rangle^2/(\lVert\mathbf{S}_a\rVert_F^2\lVert\mathbf{S}_b\rVert_F^2)$ | ∝ ε |
| **BW (ours)** | $2P-2(\sigma_1+\sigma_2)$ | constant |

### 2.4. The rank-two Bures–Wasserstein distance

To compare two pairs of weighted eigenvectors while respecting the symmetries of **P**_*t*_, we align them using an orthogonal Procrustes transform [10]. Let 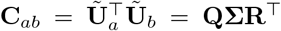 be the singular value decomposition with **Σ** = diag(*σ*_1_, *σ*_2_). The Procrustes transform **Ω** = **RQ**^*⊤*^ rotates or mirrors **Ũ** _*b*_ to best match **Ũ** _*a*_, and the remaining distance is

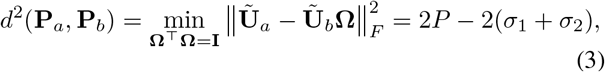

where 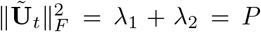. The rotation corresponds to the global oscillation phase, while the reflection reverses all lags, exactly the operations that leave **P**_*t*_ unchanged. Since the rows of **Ũ**_*t*_ are phasors, (3) gives the total squared displacement of the channel phasors after the best common rotation (Fig. 1e), measuring the phase shift required to transform one iPL matrix into the other. The square root of (3) is the Bures–Wasserstein distance between the positive semidefinite matrices **P**_*a*_ and **P**_*b*_ [11, 12]; *i*.*e*., *the optimal-transport (2-Wasserstein) distance between zero-mean Gaussian distributions with these covariances, here restricted to rank two. Since the singular values of* ***C***_*ab*_ equal those of [**c**_*a*_, **s**_*a*_]^*⊤*^[**c**_*b*_, **s**_*b*_], no eigendecomposition is required.

The BW distance combines the cosine and sine overlaps used separately by the methods in Table 1, and locally it is proportional to phase differences (extrinsic dissimilarity): for the example above, 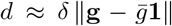, independent of *ε*. It is the only distance in Table 1 that captures the full phase pattern and measures phase differences rather than just cosine differences.

### 2.5. Bures–Wasserstein *K*-means

To identify recurring states, we combine the BW distance with *K*-means. Each state *k* is represented by a *P*× 2 centroid **M**_*k*_. Iterations alternate between (i) assigning each time point to the state minimizing 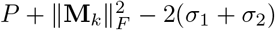, where *σ* are the singular values of 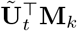, and (ii) updating each centroid by averaging its members after Procrustes alignment to the current centroid, which is one sweep of generalized Procrustes analysis [16]. We used *K*-means++ initialization [17] with five restarts. Like the data, each state 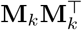 has two eigenvectors and two eigenvalues.

## 3. EXPERIMENTS AND RESULTS

### 3.1. Simulations

We compared BW-enabled *K*-means with *K*-means in the geometry of each distance listed in Table 1, using diametrical clustering [5] for the sign-invariant LEiDA and sine variants. All methods used identical initialization and five restarts. We simulated four states on the 64-channel montage of Sect. 3.2: anterior–posterior (AP) and left–right (LR) phase gradients, each with small and large lags (Fig. 2, top). For each sample, we assigned a random global phase and added independent Gaussian phase noise with standard deviation *σ* to each channel, generating 250 samples per state and repeating the experiment 20 times per noise level. Clustering accuracy was measured by normalized mutual information (NMI) between the inferred and the true states. BW *K*-means recovered the four states almost perfectly up to moderate noise and was the most accurate method at all noise levels (Fig. 2). Grassmann and sine clustering separated the two gradient orientations but merged small and large lags, even without noise, as predicted in Sect. 2.3. LEiDA, EiDA, and Frobenius separated all states at low noise but degraded considerably faster than BW. When states differed only in spatial pattern, BW performed on par with Grassmann and sine clustering and again outperformed LEiDA, EiDA, and Frobenius.

**Fig 2.**
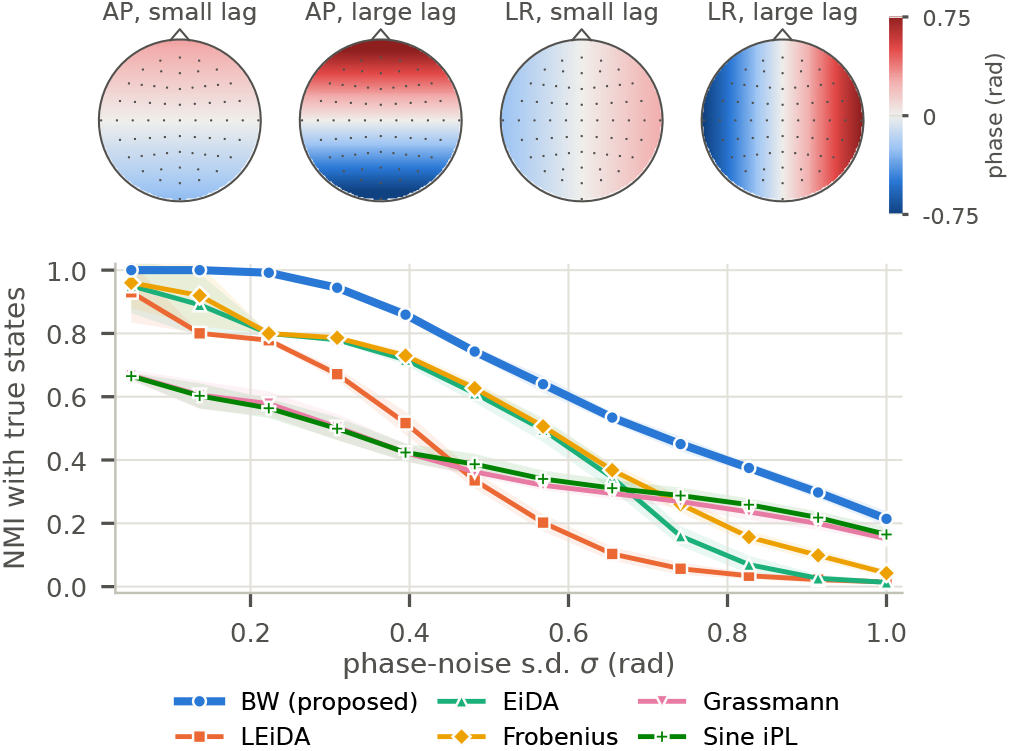
Simulations. Top: relative phase of the four true states. Bottom: NMI between estimated and true state assignments (mean ± s.d. over 20 repetitions).

### 3.2. Resting-state EEG

We analyzed 1-min eyes-closed resting-state runs (64 channels, 160 Hz) of 109 subjects in the PhysioNet EEG Motor Movement/Imagery dataset [18, 19]. Signals were filtered to 8–13 Hz (third-order Butterworth, zero phase). Each method was fitted with *K* = 4 states, as typically used for microstates [3], using 100 random time points per subject and run; all samples were then assigned to the nearest state.

The methods partitioned the data differently beyond the global state (Fig. 3). LEiDA assigned 52% of samples to a synchronous global state, while its remaining states were large-lag patterns separating channels into two nearly antiphase groups, as expected from the sign structure of the leading eigenvector. Grassmann clustering identified gradients of different orientations but no near-synchronous state; each of its three largest states mixed near-synchronous and similarly oriented lagged samples, with about 40% near-synchronous according to BW. BW *K*-means separated a near-synchronous state from an LR phase gradient and small- and large-lag AP patterns. About one-third of LEiDA’s global-state samples belonged to BW’s gradient states, indicating that the global state conceals graded lags revealed by the second, eigenvalue-weighted eigenvector.

**Fig 3.**
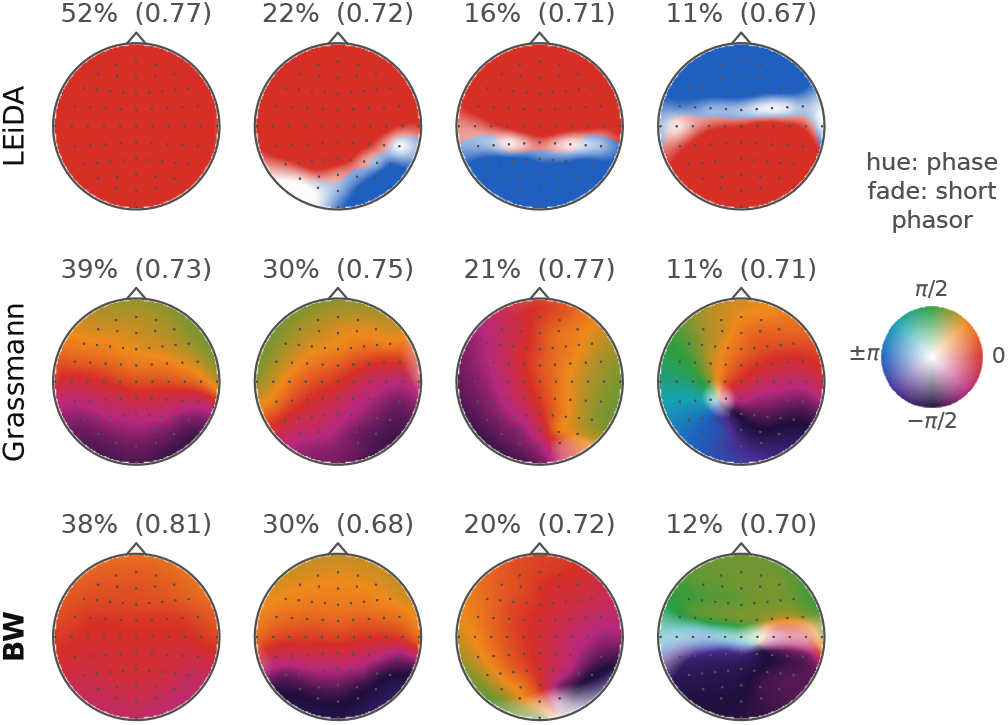
Resting-state EEG phase-locking states (*K* = 4, 8– 13 Hz, 109 subjects). Maps show the relative phase of each state; i.e., the arg of rows of **M**_*k*_. LEiDA encodes signs as phases of 0 or *π*, and Grassmann weights both columns equally.

## 4. DISCUSSION AND CONCLUSION

We showed that, for real-valued iPL matrices, comparing eigenvalue-weighted eigenvector pairs with an orthogonal Procrustes transform yields the rank-two BW distance, which measures the phase displacement between time points. Existing distances either discard information, as in LEiDA, or compare cosine and sine overlaps that become insensitive to lag size near synchrony, the typical EEG regime. Accordingly, BW was the most accurate in simulations and separated graded phase lags in EEG that LEiDA merged into a global state and Grassmann mixed with near-synchronous samples. This suggests that LEiDA’s global state may partly arise from its representation. Because the sine iPL is determined by the two eigenpairs and the BW distance combines cosine and sine overlaps, discarding the cosine to avoid volume conduction [9] is unnecessary to access lag structure. Unlike microstates, which cluster polarity-invariant amplitude topographies, BW states describe graded lags consistent with traveling waves [20], which microstates can represent only as sequences of maps. These conclusions hold for the real-valued iPL matrix; the complex-valued phase coherence matrix 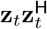 has a different geometry [6].

## 5. ETHICS AND ACKNOWLEDGEMENTS

This retrospective study used publicly available human-subject data from PhysioNet [18, 19]. The authors declare no relevant financial or non-financial interests. Claude (Anthropic) assisted with numerical verification of the derivations, analysis code, and figures; all content was reviewed and verified by the authors. Code: https://github.com/MlhMri/Clustering-iPL-states.

## Notes

### Competing Interest Statement

The authors have declared no competing interest.

